# Murine Parainfluenza Virus Persists in Lung Innate Immune Cells Sustaining Chronic Lung Pathology

**DOI:** 10.1101/2023.11.07.566103

**Authors:** Ítalo Araujo Castro, Yanling Yang, Victoria Gnazzo, Do-Hyun Kim, Steven J. Van Dyken, Carolina B. López

## Abstract

Respiratory viruses including the human parainfluenza viruses (hPIVs) are a constant burden to human health, with morbidity and mortality frequently increased after the acute phase of the infection. Although is proven that respiratory viruses can persist *in vitro*, the mechanisms of virus or viral products persistence, their sources, and their impact on chronic respiratory diseases *in vivo* are unknown. Here, we used Sendai virus (SeV) to model hPIV infection in mice and test whether virus persistence associates with the development of chronic lung disease. Following SeV infection, virus products were detected in lung macrophages, type 2 innate lymphoid cells (ILC2s) and dendritic cells for several weeks after the infectious virus was cleared. Cells containing viral protein showed strong upregulation of antiviral and type 2 inflammation-related genes that associate with the development of chronic post-viral lung diseases, including asthma. Lineage tracing of infected cells or cells derived from infected cells suggests that distinct functional groups of cells contribute to the chronic pathology. Importantly, targeted ablation of infected cells or those derived from infected cells significantly ameliorated chronic lung disease. Overall, we identified persistent infection of innate immune cells as a critical factor in the progression from acute to chronic post viral respiratory disease.

## Introduction

Infections with respiratory RNA viruses pose a constant threat to human health. It is estimated that Respiratory Syncytial Virus (RSV) alone is associated with more than 149,000 fatal cases of lower respiratory tract infection worldwide every year^1^. Among paramyxoviruses, human metapneumovirus (hMPV) is responsible for approximately 643,000 hospitalizations and >16,000 deaths globally^2^ every year, while hPIV accounts for 725,000 hospitalizations and >34,000 deaths^2,3^. Within the pediatric population, acute lower respiratory infections of viral etiology remain the leading cause of mortality in the absence of a pandemic^4,5^.

In addition to the public health burden, acute respiratory viral infections at an early age are associated with the development of chronic lung diseases including asthma and chronic obstructive pulmonary disease (COPD), while infections later in life can lead to severe exacerbations of these conditions^6,7^. Influenza, RSV, hMPV, hPIV, and rhinovirus infections have been linked to development and progression of COPD and lung fibrosis in humans, and more recently SARS-CoV-2 infection was implicated in chronic lung diseases^8–10^.

It has been long established that RNA viruses can persist in humans. The best studied example is measles virus that persists in the central nervous system of patients with subacute sclerosing panencephalitis^11,12^. Other examples include Ebola virus persistence in the testis^13^ and chikungunya virus persistence in the joints^14^. In addition, accumulating evidence suggests that respiratory viruses can establish persistent infections in the lung^15–17^. A comprehensive screening of post-mortem tissue from fatal COVID-19 cases indicated presence of viral proteins, viral RNA, and infectious SARS-CoV-2 in the respiratory tract and other anatomical sites for longer than 30 days after symptom onset^15^. Prolonged viral shedding has been reported in stem cell transplant recipient patients^18^ and immunocompromised patients infected with hPIV^16,18^. In addition, high detection rates of hPIV3 in turbinate epithelial cells were reported in patients suffering from post-viral olfactory dysfunction^19^. Prolonged exposure to viral RNA and antigens, even in the absence of infectious viral particles, can work as immune stimulation factors and contribute to chronic inflammation^20,21^. Therefore, it is critical to better understand the mechanisms of virus or viral products persistence, the sources of the virus, and their impact on chronic respiratory diseases.

Here, we used respiratory infection with the murine paramyxovirus Sendai (SeV; recently renamed Murine respirovirus) to model hPIV infections in mice and study the persistence of virus and viral products and their impact on chronic lung disease. We show that viral antigens and RNA are present in specific innate immune cells populations in the lung long after the acute infection has been cleared. Importantly, we demonstrate that infected cells, as well as cells derived from infected cells, play critical roles in maintaining post-viral chronic lung pathology.

## Results

### Severe SeV lower respiratory tract infection leads to long term persistence of viral protein and RNA in the lung

To evaluate whether mouse parainfluenza viruses persist in the respiratory tract, we infected mice intranasally with a sublethal dose of SeV strain 52 that induced severe respiratory disease, as we and others have described previously^22,23^. We monitored disease progression from 3 to 49 days post-infection (dpi), analyzing virus load and presence of virus proteins in the lungs at these time points (**Figure 1A**). Day 49 post infection has been established as a standard timepoint to study post-SeV chronic lung disease as pathology has plateaued by then^24^. Acute weight loss following SeV infection was maximal between days 7 and 8 post-infection, with animals losing about 25% of their original weight (**Figure 1B**). All mice fully recovered their weights by day 35 post-infection. As expected, virus RNA and infectious particles were detected in whole lung homogenates on day 3 post-infection, however, only viral RNA was detected on day 49 post-infection (**Figure 1C**). Immunofluorescence analysis for SeV nucleoprotein (NP) and RNAscope analysis for NP RNA (mRNA and genomic RNA) revealed a different distribution of NP in the acute and chronic phases of the infection. While NP was mostly detected in the airways lining epithelium on 3 dpi, SeV positive signal on day 49 post-infection was mostly found in the alveolar compartment associated with infiltrating cells (**Figure 1D**).

**Figure 1.**
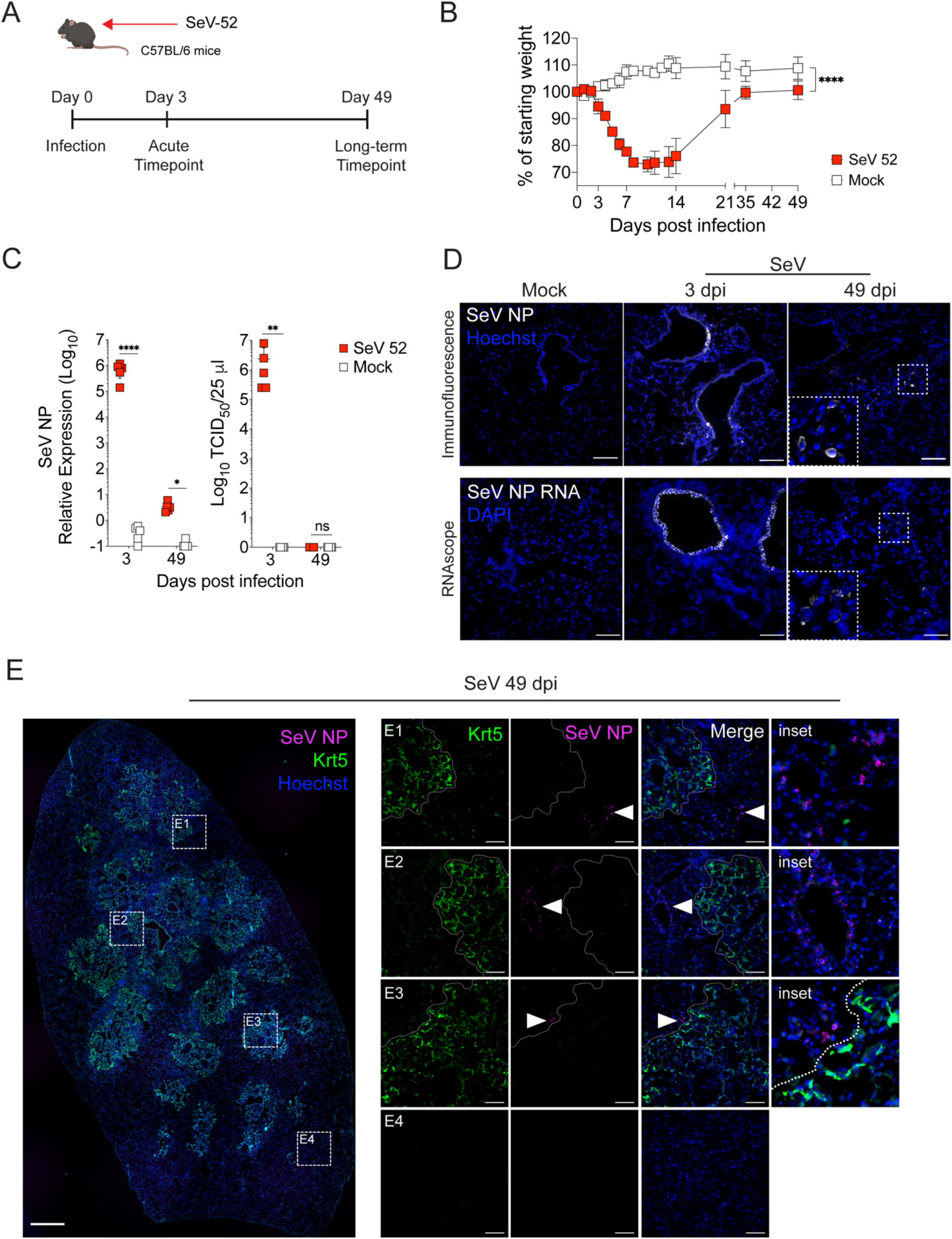
Viral antigens and RNA persist in mouse lungs after SeV-driven acute illness. **A.** Timeline of the study design. Mice were inoculated with either phosphate-buffered saline (mock) or 5×10^4^ tissue culture infectious dose (TCID_50_) of SeV 52 per animal. Lungs were analyzed on days 3 and 49 post-infection. **B.** Disease progression was monitored by measuring weight loss through the experiment. Data are representative of 4 independent experiments (mean ±SD). For SeV-infected and mock groups the area under the curve (AUC) was calculated and t-tests were performed for statistical significance analysis. ****P<0.0001. **C.** Whole lung homogenates were harvested at days 3 and 49 post-infection and both SeV NP RNA expression and infectious virus titers were quantified by qPCR and infectivity assays respectively. Relative RNA quantitation by qPCR was normalized to mouse GAPDH and β-Actin. Two-way analysis of variance with Holm-Sídák post-test was used to estimate statistical significance between groups. N = 5 animals per group. *P<0.05; **P<0.01; ****P<0.0001. **D.** SeV-infected lungs were stained for SeV NP (white staining, upper panels) using immunofluorescence and for SeV NP RNA using RNAscope (white staining, lower panels). Nuclear staining (Hoechst for immunofluorescence and DAPI for RNAscope) in blue. Representative images of 3 independent experiments. Scale bars: 100 µm. **E.** SeV-infected lungs were stained for basal stem cells (Krt5^+^, green staining) and SeV NP (magenta staining) to localize SeV NP^+^ cells in relation to areas displaying chronic lesions (dashed areas, subpanels E1-E3) and unaffected areas (subpanel E4). Arrowheads indicate SeV NP^+^ cells, more detailed in the correspondent zoomed inset panels. Images were taken using a widefield microscope. Left panel, tiling image, 5x magnification. Right subpanels, 20x magnification. Right insets: digital zooms from the correspondent 20x magnification images. Scale bars: Left panel. 500 µm; Subpanels. 100 µm. Images are representative of 3 independent experiments, 5 mice per group.

The chronic pulmonary disease caused by SeV was marked by intense tissue remodeling, with expansion of basal epithelial cells (Krt5^+^) in well-delimited lesions. It was noticeable that cells persistently expressing viral NP were adjacent to Krt5^+^ lesions, frequently organized in patches (**Figure 1E, subpanels E1-3**), while no NP^+^ cells were found in the unaffected areas of the lung (**Figure 1E, subpanel E4**). Overall, these observations demonstrate that viral RNA and viral proteins are detectable in the chronic phase of the infection, weeks after infectious virus has been cleared^23^, and indicate that there is a differential distribution of virus-infected cells in the lung during the acute (day 3) and chronic (day 49) phases of the infection.

### Persistent SeV proteins are detected in type 2 innate lymphoid cells (ILC2s), macrophages and dendritic cells

Given the diversity of cells that compose the lung inflammatory microenvironment during SeV-driven chronic disease and that most of the of persistent SeV NP^+^ cells were in the alveolar compartment (**Figure 1D, E**), we hypothesized that the persistent NP signal was mostly related to resident and infiltrating immune cells. We then used a panel of markers representing cell types frequently found in SeV-driven infiltrates in mouse lungs to characterize SeV NP^+^ cells by immunofluorescence and flow cytometry. Initially, after using antibodies against CD3 (T lymphocytes), CD11c and CD11b (dendritic cells -DCs), F4/80 (macrophages) and Thy1.2 (T cells and ILCs), we observed that CD11c^+^, CD11c^+^CD11b^+^, F4/80^+^ and CD3^-^Thy1.2^+^ cells were found to be positive for SeV NP in lung sections from 49 dpi (**Figure 2A**). Myeloid- cells, CD11c^+^, CD11b^+^ and double-positive cells, likely DCs (**Figure 2A-a1)** and macrophages (**Figure 2A-a2**), were found frequently co-expressing SeV NP, either diffusely dispersed through the lung section or clustered surrounding blood and lymphatic vessels. Unexpectedly, SeV NP signal was also detected in CD3^-^Thy1.2^+^ cells in a considerably high frequency (**Figure 2A-a3)**.

**Figure 2.**
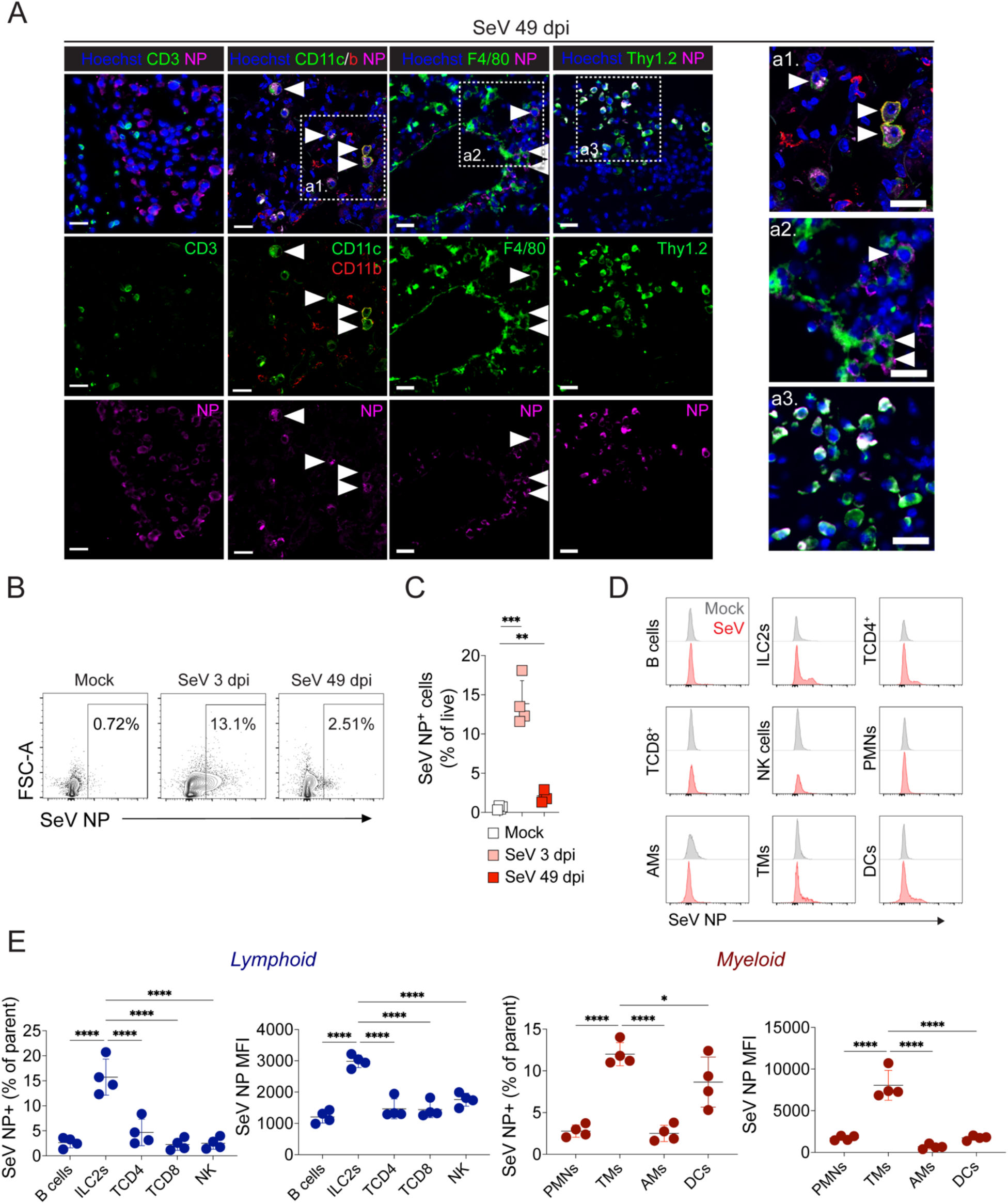
Diverse lung immune cells express SeV NP during chronic infection. **A.** Characterization of immune cells expressing SeV NP from cryopreserved mouse lungs after 49 dpi by immunofluorescence. Tissue sections were stained for SeV NP (magenta) in combination with the surface markers CD3 (T lymphocytes), CD11c, CD11b (dendritic cell subsets), F4/80 (macrophages), and Thy1.2 (Innate Lymphoid Cells and some T lymphocyte subsets) in green and red. Nuclear staining is displayed in blue. White arrows indicate individual SeV NP^+^ cells. Images were taken in a widefield fluorescence microscope using a 20x magnification scope. Scale bars: 25 µm. Representative images from three independent experiments, 5 mice per condition. Right panels: insets from the dashed areas. **B-E.** Lungs from SeV-infected mice were harvested at 3 and 49 dpi, enzymatically digested, and analyzed by multiplex spectral flow cytometry with a panel of 16 antibodies to quantify (**B and C**) and characterize (**D and E**) SeV^+^ cells. **B.** Representative dot plots of SeV NP^+^ cells (% of live) comparing acute (3 dpi) with long-term (49 dpi) SeV infection. SeV NP^+^ gates were drawn based on the isotype control and the mock-infected samples. **C.** Frequency of SeV NP^+^ cells gated on total live cells from SeV 3 days-, SeV 49 days-, and mock-infected lungs. **D.** Representative histograms of SeV NP^+^ fluorescence intensity from 9 individual cell subsets, B cells, ILC2s, T CD4^+^ lymphocytes, T CD8^+^ lymphocytes, NK cells, Polymorphonuclear cells (PMNs), Alveolar macrophages (AMs), Tissue macrophages (TMs), and Dendritic cells (DCs) at 49 dpi. Histograms from SeV-infected animals are displayed in red while histograms from mock-infected animals are displayed in gray. **E.** Frequency and mean fluorescence intensity (MFI) of SeV NP^+^ cells within lymphoid- and myeloid-origin cell subsets. All multiple comparisons were done with one-way ANOVA and Holm-Sídák post-test. *P<0.05; **P<0.01; ***P<0.0005; ****P<0.0001. Data representative of two independent experiments, 4-5 animals per condition, total 1 million events acquired per animal.

**Figure 3.**
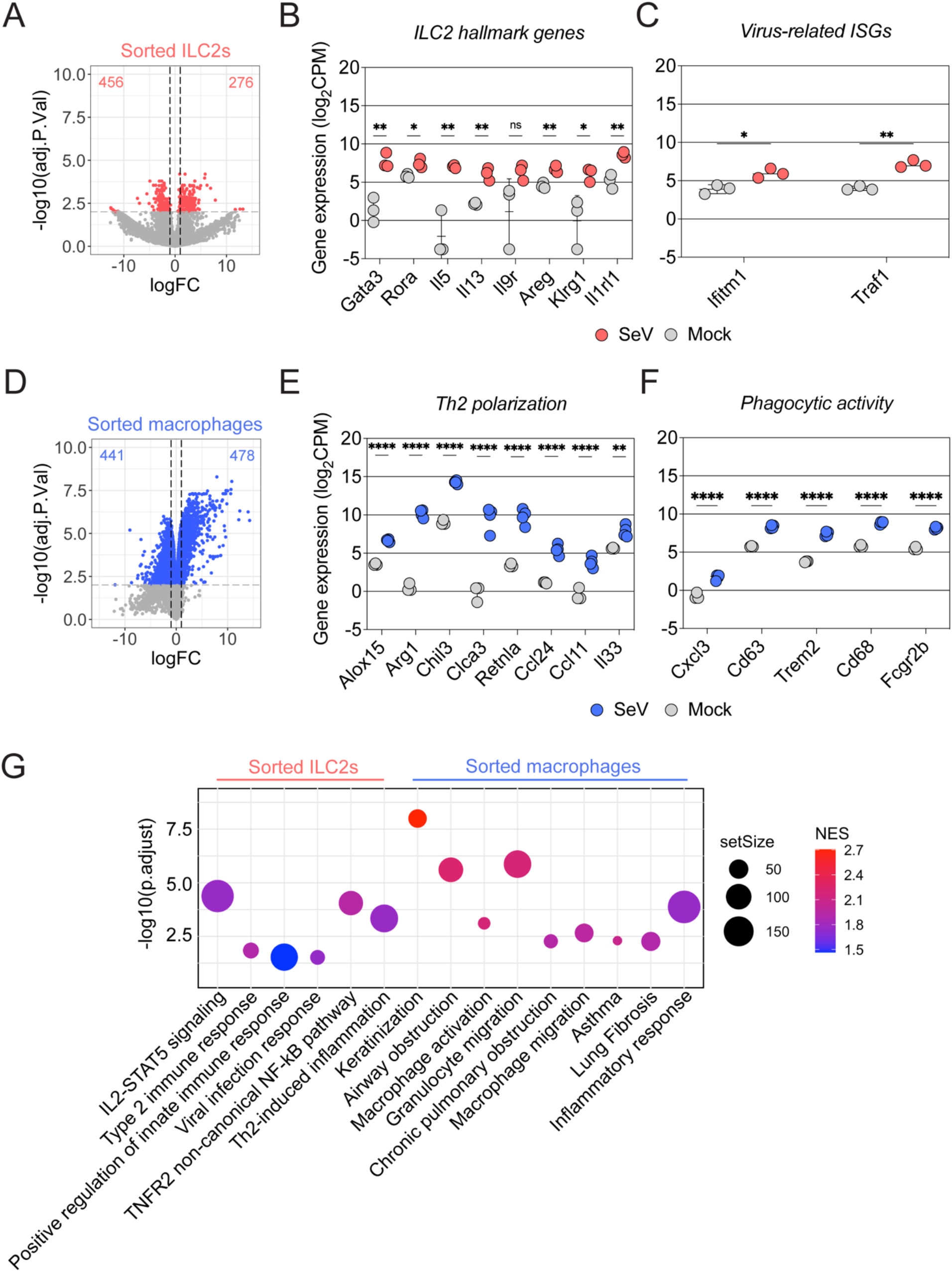
Type 2 innate lymphoid cells and macrophages are persistently activated in a type 2 inflammation manner during SeV chronic lung disease. Lung type 2 innate lymphoid cells (ILC2s) (**A-C**), and macrophages (**D-F**) were isolated either from SeV- or mock-infected mouse lungs after 49 dpi and subjected to bulk RNA-seq. **A** and **D.** Volcano plots indicating differentially expressed genes detected in ILC2s and macrophages, respectively, from SeV-infected lungs over mock. P<0.05, LogFC>2. **B-C.** Scattered dot plots showing expression of ILC2 hallmark genes (**B**) and virus-related ISGs (**C**). Each dot corresponds to an individual pool of cells (n = 6 animals pooled in pairs per condition). **E-F.** Scattered dot plots indicating expression of Th2 polarization (**E**) and phagocytic activity (**F**) genes from macrophages. Each dot corresponds to cells obtained from an individual animal (minimum n = 3 animals per condition). Data are displayed as mean ± SD. Two-way analysis of variance (ANOVA) with Bonferroni post-test was used to estimate statistical significance between multiple comparisons. *P<0.05; **P<0.01; ****P<0.0001. CPM, copies per million. **G.** Bubble chart showing gene set enrichment analysis (GSEA) of upregulated genes in ILC2s and macrophages sorted from SeV 49 dpi lungs. Bubble size indicates gene set size per GSEA pathway, while bubble color gradient indicates Normalized Enrichment Scores (NES) values.

To quantify and better characterize the cell subtypes that are sources of persistent virus antigens, we employed multiplex spectrum flow cytometry to analyze SeV-infected mouse lungs at the chronic stage of the infection (**Figure S1**). Single-cell suspensions obtained from acute SeV infection yielded on average 13.8±2.9% live, NP^+^ cells. On day 49 after infection, the percentage of live NP^+^ cells was 2.12±0.7% (representative dot plots in **Figure 2B** and quantitative analysis in **Figure 2C**). We then analyzed individual cell subtypes to determine the percentage of SeV NP^+^ cells withing each subset. Amongst lymphoid-origin cells, ILC2s displayed the highest percentages and MFI of NP expression (15.7±3.5%) (**Figure 2D** and **E**, left panels). Meanwhile, tissue macrophages (TMs) (12.01±1.3%) and DCs (8.65±3%) were the myeloid subsets with the highest percentages of NP signal among the cell types analyzed (**Figure 2D** and **E**, right panels). Altogether, our data strongly indicate that macrophages, ILC2s and dendritic cells are the main sources of persistent virus antigens during chronic SeV infection.

### Innate immune cells sustain type 2-inflammation during SeV-driven chronic lung disease

Macrophages, dendritic cells and ILC2s participate in the establishment of chronic lung disease upon SeV infection through the IL-33/IL-13 axis^25^. Finding these cells harboring persistent viral products raised additional questions about their impact on the pathogenesis of chronic lung disease. We sorted ILC2s and macrophages from SeV-infected IL-13 reporter (sm13)^26,27^ mouse lungs at 49 dpi and analyzed their transcriptomic signatures against mock-infected animals to characterize their activation state. ILC2s displayed a total of 732 differentially expressed genes (DEGs) over mock, with 276 being upregulated and 456 being downregulated (P<0.05 and LogFC>2) (**Figure 3A**). We observed a significant increase of ILC2 hallmark genes^28^, including *Gata3*, *Rora*, *IL5*, *IL13*, *Areg*, *Klrg1*, and *Il1rl1* (ST2) (**Figure 3B**), as well as genes known to be increased in the context of RNA virus infection (**Figure 3C**). Gene set enrichment analysis (GSEA) showed that lung ILC2s from SeV-infected were enriched in pathways previously linked to SeV infection^29^, including the TNFR2 non-canonical NF-kB pathway (**Figure 3G**). Moreover, DEGs from sorted ILC2s show signatures associated with type 2 immune responses and inflammation, supporting the role of ILC2s in maintaining the chronic type 2 immunopathology observed in SeV-infected lungs at 49 dpi (**Figure 3G**). Lung macrophages from SeV-infected mice displayed 478 upregulated DEGs compared to mock (**Figure 3D**), from which classical genes involved in Th2 inflammation^30^, including *Arg1*, *Chil3*, *Ccl11*, *Il1rl1* (ST2), and *Il33* were significantly increased (**Figure 3E**). In addition, these macrophages had transcriptomic signatures associated to increased phagocytic activity^31^ such as *Cd63*, *Cd68*, and *Fcgr2b* (**Figure 3F**). Similar to what was shown for ILC2s, upregulated genes in macrophages were enriched in GSEA signatures directly involved in chronic lung disease (**Figure 3G**). These findings indicated that innate immune cells display a strong gene expression polarization towards type 2 inflammatory responses and show transcriptomic footprints that are indicative of virus infection and chronic lung diseases, confirming their involvement in the pathogenesis of post-viral chronic lung disease.

### Virus-infected cells and/or cells derived from them contribute to the chronic inflammatory state following SeV infection

Long-term exposure to viral antigens and RNA in the respiratory tract is likely to have important implications on the tissue microenvironment. To directly evaluate the transcriptome of persistently infected cells, we generated a double-reporter virus by inserting an eGFP and a Cre recombinase as independent genes between the N and the P genes of the SeV Cantell strain (rSeV-C^eGFP-Cre^) (**Figure S2A**). We then used rSeV-C^eGFP-Cre^ to infect tdTomato reporter mice that contain a Cre reporter allele flanked by loxP-STOP cassettes (**Figure S2B**). We harvested lungs on day 49 pi for flow cytometry and FACS analysis (**Figure 4A**). Infection with rSeV-C^eGFP-Cre^ progressed in a similar way as our previously described SeV-52 model. Weight loss of up to 20% of the original weight was observed until day 9 post-infection, with steady recovery until no differences from uninfected animals by day 21 (**Figure 4A**). Upon transcription of the viral genome, Cre is expressed and excises the loxed stop cassette that blocks the tdTomato gene construct leading to expression of the reporter. Virus-infected cells, cells that cleared the infection in a non-cytolytic manner, and cells derived from these cells will have constitutive expression of the tdTomato fluorescent protein (tdTom) (**Figure S2B**). At 3 dpi, the frequency of tdTom^+^ cells in infected lungs was 21.3%±0.6%, and this number decreased to 10±2.6% at 49 dpi (**Figure 4B-C**, includes a representative sample). From these cells, at 3 dpi 84.4%±3.8% were characterized as non-immune (CD45^-^) and 15.4%±3.8% were characterized as immune cells (CD45^+^) (**Figure 4D-E**, includes a representative sample), similar to what we observe during SeV-52 acute infections. The proportion of non-immune tdTom^+^ cells at 49 dpi remained high (90.9%±0.3%) and immune cells accounted for 8.8%±0.2% of all tdTom^+^ events (**Figure 4D-E**, includes a representative sample). As previously reported, the constitutive expression of tdTomato upon Cre exposure is independent of further viral replication, and all tdTom^+^ cells, even those that cleared the infection would express the reporter protein^32^. To differentiate cells persistently expressing viral protein from all other tdTom^+^ cells, we combined tdTom detection with SeV NP staining. Using this strategy, we identified three distinct cell populations in tdTom reporter mice on day 49 after infection with rSeV-C^eGFP-Cre^: tdTom^+^NP^-^, tdTom^+^NP^+^, and negative cells (**Figure 4F**). We then sorted these populations for transcriptome analysis. To increase RNA yields, we used three pools of two individual infected animals for sorting. In addition, we obtained negative cells from mock-infected tdTom reporter animals as controls. The samples were subjected to total RNA-seq. We found reads mapping to the viral genome or the eGFP reporter gene in all three tdTom^+^NP^-^ cell pools and in all three tdTom^+^NP^+^ pools (**Figure 4G**). Alignment to the rSeV-C^eGFP-Cre^ genome showed viral reads mapping to multiple viral genes, including the polymerase L gene in some of the pools (**Figure H**), confirming that tdTom^+^NP^-^ cells have been exposed to the virus either by direct infection or through their progenitors.

**Figure 4.**
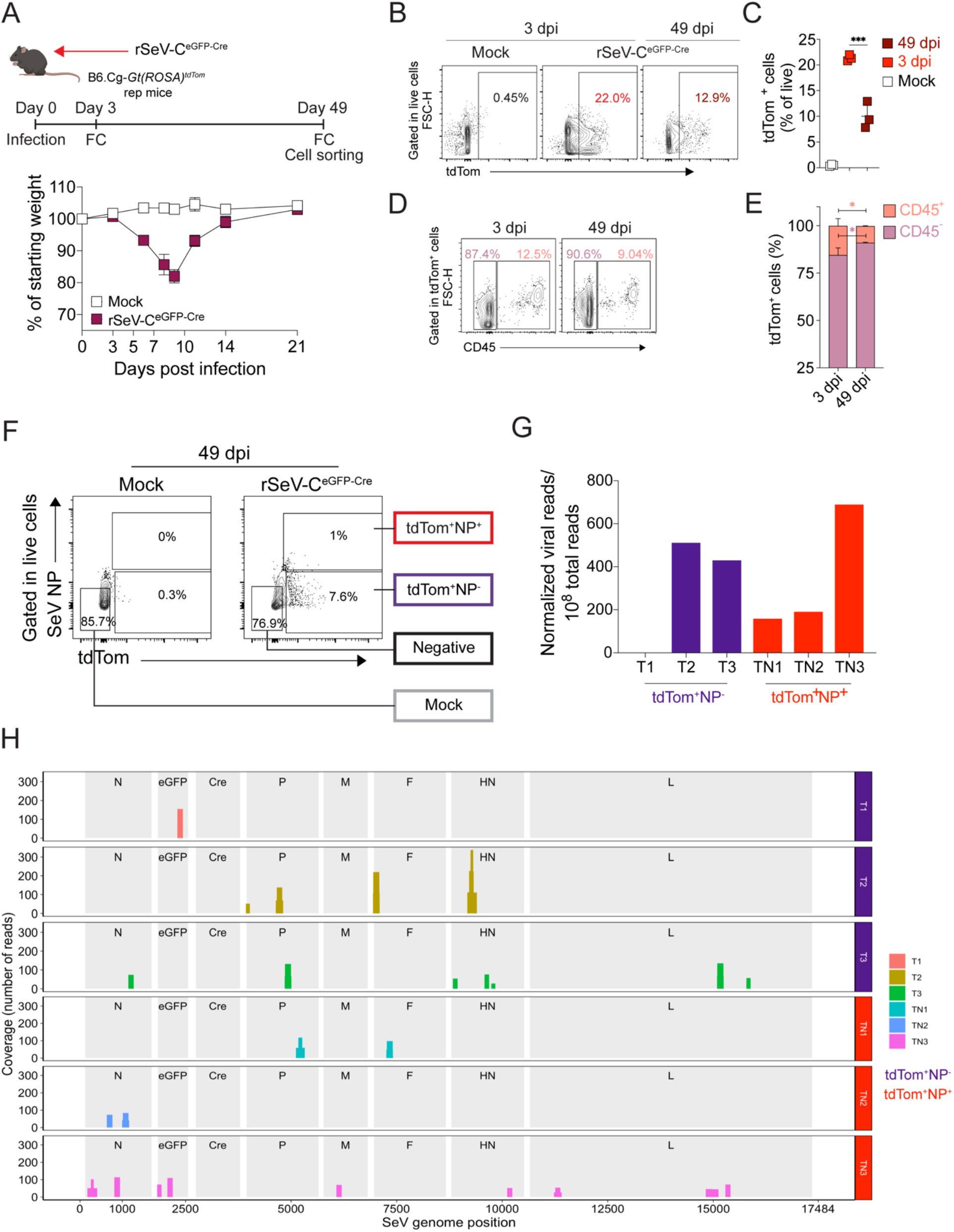
Paramyxovirus infection clearance is not complete and leaves long-term survivor cells in the lower respiratory tract expressing persistent viral RNA and viral proteins. **A.** B6.Cg-*Gt(ROSA)*^tdTom^ (tdTom) mice were inoculated intranasally with either PBS (mock) or 5×10^5^ TCID_50_ rSeV-C^eGFP-Cre^. Lungs were harvested at 3, and 49 dpi for flow cytometry (FC) analysis, and cell sorting at 49 dpi. Weight loss was recorded up to 21 dpi to monitor disease progression. Data (mean±SD) are representative of 2 individual experiments (minimum 3 mice per group). **B.** Representative dot plots comparing percentage of tdTom^+^ cells in mouse lungs during acute (3 dpi) and chronic (49 dpi) rSeV- C^eGFP-Cre^ infection. **C.** Frequency of tdTom^+^ cells gated on total live cells from rSeV-C^eGFP-Cre^ 3 days-, rSeV- C^eGFP-Cre^ 49 days-, and mock-infected lungs. Data are shown as mean±SD. Statistical significance was estimated with one-way ANOVA using Bonferroni post-test. ***P<0.005. **D-E**. Characterization of immune (CD45^+^) and non-immune (CD45^-^) cell proportions within tdTom^+^ cells. Representative dot plots (**D**) and quantification (**E**) of CD45 staining in tdTom^+^ cells during acute (3 dpi) and chronic (49 dpi) rSeV-C ^eGFP-Cre^ infection. Statistical significance was estimated with two-way ANOVA and Holm-Sídák post-test. *P<0.05. **F.** Combination of tdTom and SeV NP detection enables sorting of two distinct subsets of VID cells, SeV-infected cells persistently expressing viral antigens (tdTom^+^NP^+^) and SeV-infected/survivor cells only (tdTom^+^NP^-^). Representative dot plots indicating the gating strategy used for sorting 3 cell subpopulations from rSeV-C^eGFP-Cre^-infected lungs tdTom^+^NP^-^, tdTom^+^NP^+^, and negative. Negative cells from mock-infected lungs were also sorted. Data representative of 2 individual experiments, 6 mice per condition. **G.** Sorted cells were pooled (2 mice) and subjected to RNAseq. Normalized viral reads per 10^8^ total reads are displayed per individual pool of tdTom^+^NP^-^ and tdTom^+^NP^+^ cells. **H.** Coverage analysis indicating normalized viral reads per genome position in each individual cell pool. Viral specific gene regions and reporter genes are indicated in light gray.

After identifying two major cell subsets relevant to virus-host interactions in persistent SeV infection, we sought to understand their role in pathogenesis by comparing the host transcriptomic profiles of tdTom^+^NP^-^ and tdTom^+^NP^+^ cells. Transcriptomic analysis of the different pools showed differentially expressed genes (DEGs) in tdTom^+^NP^-^ and tdTom^+^NP^+^ cells over mock (**Figure 5A**) (p<0.05 and LogFC>2). Negative cells showed no significant transcriptomic changes over mock (**Figure 5A**), suggesting that only direct virus-host interactions lead to long-term host gene expression changes, rather than a more widespread effect to non-infected cells in the lung.

**Figure 5.**
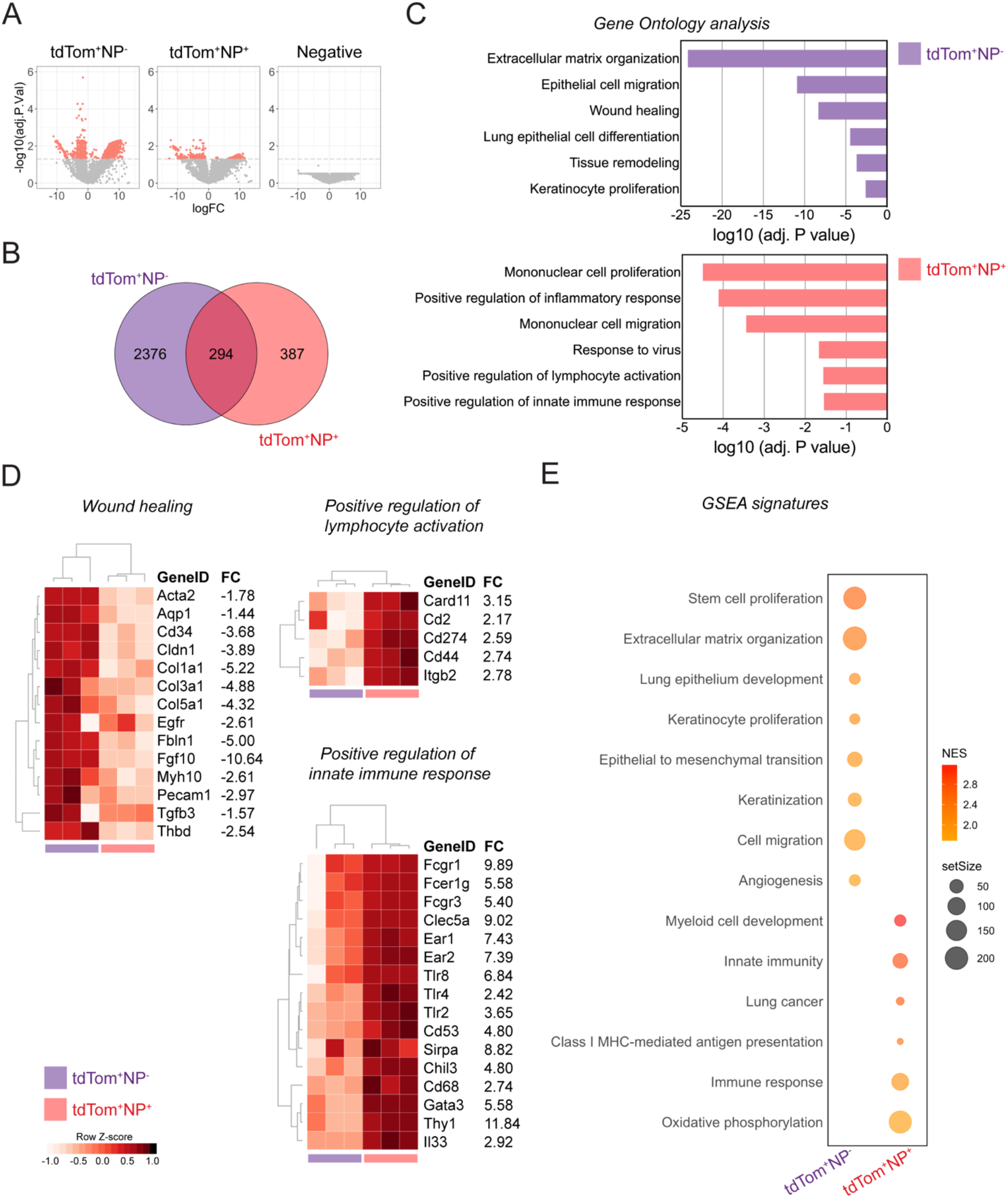
Viral clearance and persistence entails opposing transcriptional programs in long-term SeV-infected lungs. Viral infected and cells derived from them sorted from tdTom mice infected with rSeV-C^eGFPCre^ at 49 dpi were subjected to bulk RNAseq and host transcriptome analysis. **A.** Volcano plots indicating differentially expressed genes (DEGs) in tdTom^+^NP^-^, tdTom^+^NP^+^, and Negative cells against Mock negative cells. **B.** Venn diagram showing overlapping DEGs from tdTom^+^NP^-^ and tdTom^+^NP^+^, as well as exclusive DEGs from each cell subset. **C.** Bar graphs showing gene ontology (GO) enrichment analysis of each of the VID cell subsets (tdTom^+^NP^-^ and tdTom^+^NP^+^) exclusive DEGs. **D.** Heatmaps of selected gene collections from the GO pathways in **C.** Shown are fold change (FC) values of tdTom^+^NP^-^ and tdTom^+^NP^-^ are displayed. Columns groups are color-coded following the same patterns on **C.** and **B. E.** Gene set enrichment assay (GSEA) bubble chart indicating the most significant enriched pathways of tdTom^+^NP^+^ transcriptome signatures in comparison with tdTom^+^NP^-^ cells. Bubble color gradient indicates Normalized enrichment score (NES), and bubble sizes correspond to gene set size on each pathway.

From the identified DEGs, 2376 were exclusive of tdTom^+^NP^-^ and 387 DEGs were exclusive of tdTom^+^NP^+^ cells (**Figure 5B**). Gene Ontology (GO) analysis indicated that tdTom^+^NP^-^ cells have gene expression signatures associated to extracellular matrix organization, epithelial cell migration, wound healing, lung epithelial cell differentiation, tissue remodeling and keratinocyte proliferation (**Figure 5C**), of note, the extracellular matrix organization factors *Col1a1*, *Col3a1*, *Col5a1*, and *Fbln1*, and the cell proliferation factors *Cd34*, *Tgfb3*, *Cldn1*, *Egfr*, and *Fgf10*, all included in the wound healing pathway (**Figure 5D**).

Differently, tdTom^+^NP^+^ DEGs included genes associated with mononuclear cell proliferation and migration, response to virus, positive regulation of inflammatory response, lymphocyte activation, and innate immune response (**Figure 5C**). Genes involved in lymphocyte activation and proliferation included *Cd44*, *Itgb2*, *Cd274*, *Cd2*, and *Card11*. Among genes involved in positive regulation of innate immune responses, *Cd68*, *Chil3*, *Fcgr3*, *Fcer1g* and *Fcgr1* are examples of myeloid-related genes, *Gata3*, and *Thy1* are examples of innate lymphoid-related genes (**Figure 5D**), and *Il33* is a known gene involved in Type-2 immunopathology^22^.

Gene set enrichment analysis (GSEA) comparing tdTom^+^NP^-^ and tdTom^+^NP^+^ cell transcriptomes confirmed the distinct transcriptomes of these cell populations. Survivor tdTom^+^NP^-^ cells had enriched signatures associated to tissue remodeling, keratinization, and tissue homeostasis-related pathways (**Figure 5E**). On the other hand, persistent tdTom^+^NP^+^ cells had enriched signatures mapping almost exclusively to immune and inflammation-related pathways, such as myeloid cell development, innate immunity, lung cancer, and oxidative phosphorylation (**Figure 5E**). These observations indicate that while cells that cleared SeV infection initiate and maintain a transcriptomic program to control inflammation and to resolve lung injury, cells persistently exposed to viral products have a pro-inflammatory profile with potential implications on lung pathogenesis.

### Persistent infected cells and its progeny are directly involved in the progression of chronic lung disease

We next set out to investigate the impact of persistent infected cells in the lung chronic disease progression. To do this, we combined our rSeV-C^eGFP-Cre^ with Cre-inducible Diphtheria toxin receptor (iDTR) mice^33^, where SeV-infected cells constitutively express DTR. DTR-expressing cells would then be susceptible to specific ablation following DT administration. We then intranasally injected the iDTR mice with either PBS (mock) or 2×10^5^ TCID_50_/animal of rSeV-C^eGFP-Cre^, treated with two consecutive doses of DT, and analyzed their lungs by flow cytometry at 5 dpi to check for depletion of infected cells (**Figure S3A**). The DT treatment did not impact disease progression, and both infected groups lost up to 20% of their original weight and recovered thereafter (**Figure 6B**). Flow cytometry quantification of SeV NP^+^ cells indicated that two consecutive DT doses were sufficient to deplete more than 80% of infected cells (**Figure S3B-C**) as described previously^33^. To deplete persistent infected cells, we waited until 21 dpi to avoid the acute and clearance phases of SeV infection and used the same 2-dose DT regime,

**Figure 6.**
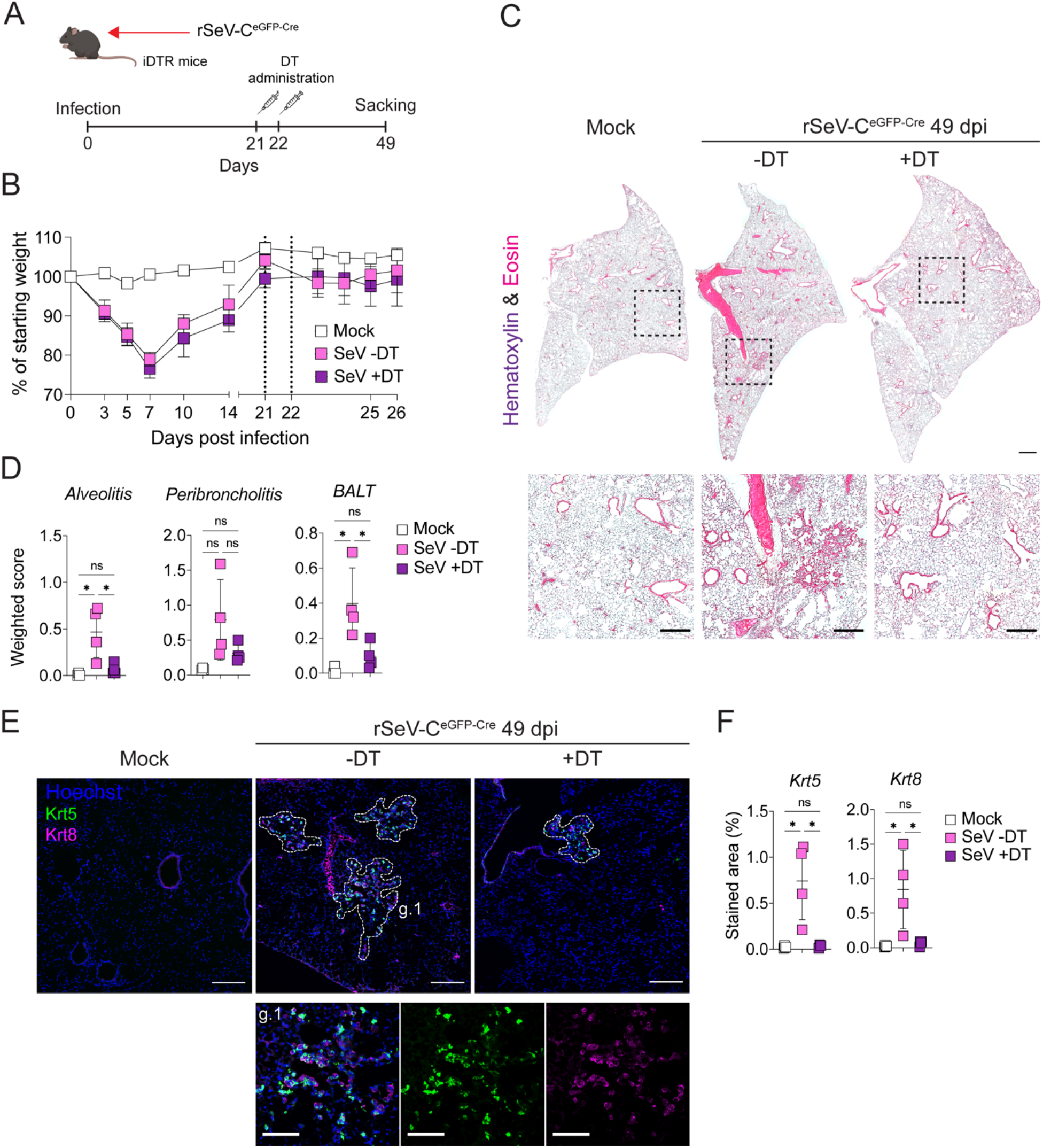
Chronic lung pathology induced by paramyxovirus infection is dependent on long-term surviving and persistently infected cells. **A.** Diphtheria toxin regime treatment and timepoints for tissue harvesting and analysis. **B.** Disease progression was assessed by monitoring animal weight loss until 26 dpi. Graphs are representative of 2 independent experiments and depict mean weight loss values ±SD, 4 mice per condition. **C.** Mouse lungs were harvested at 49 dpi and tissue sections were stained with Hematoxylin and Eosin to compare pathological changes between the analysis groups. Top panels indicate representative images of whole lung sections (Brightfield, 5x magnification tiled images, Scale bars: 1 mm) and bottom panels indicate zoomed-in images from the indicated areas (Brightfield, 5x magnification tiled images, scale bars: 100 µm). 4 animals per condition. **D.** Lung sections from SeV +DT and SeV -DT groups were blindly scored for histopathological changes. Total area affected, percentage of airway structures affected, and intensity of alveolitis, peribroncholitis, and bronchus-associated lymphoid tissue (BALT) expansion were determined for every individual lung sample. Individual weighted scores values ±SD are indicated. Data representative of 2 individual experiments, 4-7 mice per condition. **E.** Lung sections were stained for the tissue remodeling and chronic lung lesion markers Krt5 (green) and Krt8 (magenta) with immunofluorescence to check for chronic lung lesion progression. Nuclear staining is displayed in blue. The dashed area indicates chronic lung lesions and areas of intense tissue remodeling. Images were taken with a widefield microscope. Upper panels, tiling images, 20x magnification, scale bars: 500 µm. Lower panels: 20x magnification, scale bars: 100 µm. **F.** Quantification of chronic lung lesion area (%) over total lung section area. Mean values ±SD are shown. Data are representative of two individual experiments, 4-7 mice per condition. Statistical significance was estimated using one-way ANOVA and Bonferroni post-test. *P<0.05.

with the final time point for analysis at 49 dpi (**Figure 6A**). Infected groups showed the typical signs of disease progression following SeV infection, and the DT regime did not cause weight loss in the 5 days following intraperitoneal administration (**Figure 6B**). As expected, infected mice that were not treated with DT had the typical signs of chronic lung disease caused by SeV, including alveolitis, bronchiolization of the alveolar compartment, and bronchoalveolar lymphoid tissue (BALT) formation^22^ (**Figure 6C**). Noticeably, DT-mediated ablation of persistently infected and survivor cells decreased the intensity of the lung disease (**Figure 6C-D**), and lung sections displayed minimal pathology compared with untreated controls. Targeted DT ablation of SeV NP^+^ cells also significantly decreased the area affected by chronic lesions, marked by intense agglomeration of basal stem cells (Krt5^+^) and transitory epithelial cells (Krt8^+^) (**Figure 6E-F**), hence demonstrating that persistent infected cells and its progeny are directly involved in the progression of SeV-driven chronic lung disease.

## Discussion

RNA virus-host interactions are proving more complex and intricated than they were thought to be. Increasing evidence shows that a diversity of RNA viruses can remain in their host in different forms past acute illness recovery, clearance of infectious viral particles and development of specific immunity, resulting in harmful long-term clinical manifestations with epidemiological implications^20,34,35^. Frequently, persistent RNA infections are found in very specific niches, for instance, the CNS, testis, ocular tissue, and secondary lymphoid organs. Here, using a natural host-pathogen model, we describe the implications of respiratory virus persistence in a non-canonical immune privileged site, the lower respiratory tract. In accordance to previous reports for SARS-CoV2 and influenza virus^17,36^, we showed detection not only of SeV RNA from multiple viral genes for up to 49 dpi, but also of viral antigens, indicating that some level of persistent viral RNA translation is still in place during the persistent phase of the infection, even with undetectable infectious viral particles. Importantly, we identified ILC2, macrophages and dendritic cells as the main cells harboring persistent viral products in the lung, and we demonstrate that depletion of cells that have been directly infected or those derived from infected cells significantly reduce chronic post-viral lung disease.

Our data indicate that between the early stages of SeV infection and the later chronic lung inflammation there is a major shift of viral antigen and RNA cell sources (**Figures 1 and 2)**. As expected, based on previous reports^37^ airway cells were the major source of viral proteins and RNA on day 3 post-SeV infection. However, on day 49 post-infection cells in the alveolar compartment were the sources of viral products, with no detectable virus RNA nor antigens in epithelial cells. Upon immunofluorescence and flow cytometry, these viral product sources were characterized as dendritic cells, macrophages and ILC2s. One of the hallmarks of the chronic SeV-driven immunopathology is the increased number of alternatively activated macrophages (AAMs) and monocyte-derived dendritic cells in infected mouse lungs^38–40^. These cells were shown to partner with ILC2s, also found in increased numbers in this condition, contributing to chronic type 2 inflammation in an IL-33/IL-13 axis-dependent manner^25^. In light of our findings, we hypothesize that immune cells with phagocytic activity, such as macrophages and dendritic cells, interact with SeV-infected epithelial cells and become sources of viral RNA and protein, as previously shown that macrophages and dendritic cells remain positive for SeV RNA by qPCR for up to 21 dpi^40^. The transcriptomic signatures of sorted macrophages during SeV chronic lung disease support these observations, with strong Th2 polarization and increased expression of phagocytic activity-related genes (**Figure 3**), but no significant changes in markers suggestive of active viral infection (data not shown).

For the myeloid compartment, phagocytosis of viral-infected cells is a known pathway whereby professional phagocytes could acquire viral antigens and even become infected. Unexpectedly, we found ILC2s as one of the most significant sources of persistent viral subproducts following SeV infection. Since ILC2s lack phagocytic activity, our results suggest that expression of SeV antigens is a consequence of viral infection. ILC2s from SeV persistently infected lungs displayed upregulated ILC2 hallmark IL-2- STAT5 and Th2 inflammation genes, corroborating previous studies covering ILC2 roles in post-viral respiratory disease^28,41^. However, we also reported genes upregulated during viral infection, such as *Ifitm1* and *Traf1*, involved in viral infection response pathways. To the best of our knowledge, this is the first report demonstrating the persistent expression of viral antigens in innate lymphoid cells.

Typical infections caused by highly cytopathic viruses were thought to follow a canonical chain of events culminating with cell lysis, tissue damage, release of pro-inflammatory mediators, and viral clearance. Nonetheless, it was recently shown that cells can overcome viral infection in a non-lytic way and either the infected surviving cell or its daughter cells have diverse long-term implications^42–46^. For instance, influenza A virus (IAV)-directly infected or derived from infected club cells showed a pro-inflammatory profile that implicated in lung pathology in mice^42^, while also exerting a protective role against secondary infections^43^. Survivor epithelial cells from another respiratory orthomyxovirus infection, influenza B virus (IBV), were also shown to be critical to maintain respiratory barrier function in a murine model of IBV infection^45^. The infection by SeV, a murine respirovirus, also left a significant percentage of survivor cells in mouse lungs, from which the majority were epithelial cells (**Figure 4**). Interestingly, even as early as 3 dpi, CD45^+^ immune cells were positive for the reporter tdTom, indicating active viral replication or phagocytosis of infected cells. After 49 days of the infection, CD45^+^ immune cells were still part of the tdTom^+^ cell pool, suggesting that for paramyxoviruses the diversity of survivor cells goes beyond the epithelial compartment.

As suggested previously, survivor cells could act as long-term sources of viral antigens^47^ with potential role in chronic lung disease development and lung healing. We successfully combined the cell-fate tracing system using Cre-dependent tdTom reporter mice/rSeV-C^eGFP-Cre^ with virus antigen detection to better characterize survivor and virus persistent cells. For both survivor cells with or without persistent expression of SeV NP (tdTom^+^NP^-^ and tdTom^+^NP^+^), SeV RNA was detected and coverage analysis after RNAseq indicated multiple viral genes represented, including the polymerase L gene, suggestive of low replicative levels in SeV persistent cells. Based in our previous cell characterization of persistent SeV-antigen expressing cells, we hypothesized that the survivor tdTom^+^NP^+^ cells were exclusively immune cells.

Cells that manage to survive a non-cytolytic clearance of RNA viruses after the acute stages of the infection can maintain abnormal transcriptomic footprints for weeks^46^. After following SeV-infected mice for a long-term post-infection (49 dpi) but also keeping in mind that the chronic lung disease is present at this timepoint, we assessed the transcriptomic signatures of tdTom^+^ cells in comparison with negative and mock cells. Surprisingly, deep transcriptomic changes were observed only in survivor tdTom^+^NP^-^ and tdTom^+^NP^+^ cells. Negative cell transcriptomes resembled mock cells, giving no significant differentially expressed genes. A similar observation was made with uninfected cells from IBV infected mice at 14 days in comparison with their correspondent controls^45^. These findings indicate that at later stages of paramyxovirus infection, the lung transcriptomic changes are a result of direct virus-cell interaction events and not due to responses to secondary secreted mediators. Given the diversity of survivor cells that we described from a respiratory paramyxovirus infection, we expected correspondent diverse roles in pathogenesis. We observed that tdTom^+^NP^-^ cells presented gene expression signatures matching to tissue remodeling, wound healing, and lung regeneration pathways, typically seen in survival epithelial cells from respiratory orthomyxovirus infections^42,45^. However, infected cells expressing persistent SeV antigens (tdTom^+^NP^+^ cells) displayed a different gene signature enriched in proinflammatory genes, matching the gene expression profiles we described from sorted innate immune cells from SeV-driven chronic lung disease.

To address the impact of SeV-survivor cells in the subsequent chronic lung pathology, we employed Cre-inducible diphtheria toxin receptor (iDTR) mice in combination with our rSeV-C^eGFP-Cre^ virus and performed DT-mediated ablation of viral infected cells and its progeny way after the timepoints were SeV infection is considered cleared. Unfortunately, we were not able to specifically deplete persistent SeV-survivor NP^-^ or NP^+^ cell populations one at a time at this point due to lack of appropriate tools available to us. Regardless of the fact that DT ablation affects all DTR-expressing populations (persistent infected, virus-cleared, and its progeny), and not only the survivor NP^+^ cells, there was a significant impact on pathogenesis, reinforcing that the chronic Th2-biased inflammation induced by survivor SeV NP^+^ cells is a key factor to the maintenance of the “post-viral” chronic lung disease.

In summary, our study demonstrates that paramyxovirus RNA and antigens persist in the lower respiratory tract associated with innate immune cells for months after the viral infection is thought to be cleared. More importantly, clearance of SeV infection is not complete, leaving a complex collection of survivor cells including epithelial and immune cells, each carrying distinct transcriptomic signatures. Long-term expression of virus antigens by survivor cells was proven to be a key factor for the immunopathology of paramyxovirus infection as a persistent source of activation for innate immune cells, which leads to maintenance of the robust type 2 environment in the SeV-driven chronic lung disease. Our data not only shed light into the cellular fate following respiratory infections and its long-term implications on chronic disease, but also pave the road for studies to further characterize the functions of persistent virus antigen-expressing cells and designing antiviral strategies.

## Materials and Methods

### Mice, Virus Infection and Virus Titration

Seven- to 9-weeks old female wt C57BL/6 mice were either bred in house or purchased from Taconic Biosciences (Rensselaer, NY). B6.Cg-Gt(ROSA)26Sor^tm14(CAG-tdTomato)Hze^/J (tdTomato) and C57BL/6-Gt(ROSA)26Sor^tm1(HBEGF)Awai^/J (iDTR) mice were either bred in house or purchased from The Jackson Laboratory. B6.129S4(C)-Il13^tm2.1Lky^/J (sm13) mice were previously described^26,27^ and bred at Washington University in St. Louis. Sendai virus strain 52 (SeV-52) stocks were expanded in 10-days-old embryonated chicken eggs (Charles River Laboratories, Wilmington, MA) and virus titers were determined using end-point dilution tissue culture infectious dose (TCID_50_) infectivity assays^48^ in LLC-MK2 cells. For mice infections, animals were anesthetized with standard doses of Xylazine/Ketamine and injected intranasally with 40 µL of either PBS or diluted virus to a final dose of 5×10^4^ TCID_50_ for SeV-52 or 5×10^5^ TCID_50_ for rSeV-C^eGFP-Cre^, corresponding to 10 times the virus ID50 ensuring that all mice were infected. Mouse groups were then monitored for weight loss as an indicator of disease progression for up to 49 dpi. To obtain lung viral titers, lung lobes were homogenized in 0.1% Gelatin, clarified by centrifugation, and analyzed by infectivity assays in LLC-MK2 cells. iDTR mice were administered with 100 ng of Diphtheria toxin (DT) (Sigma) intraperitoneally at days 3 and 4 for depletion of rSeV-C^eGFP-Cre^-infected cells. To achieve depletion of persistently infected and survivor cells, DT treatment was performed at days 21 and 22 post-infection.

### Generation of rSeV-C^eGFP-Cre^

The full-length SeV Cantell viral antigenome sequence (NCBI accession number OR764764) was inserted into the pSL1180 vector flanked at the 5’ by the T7 polymerase promoter and a Hammer-head Ribozyme (Hh-Rbz) and at the 3’ by a second Ribozyme, and the T7 terminator. The reporter eGFP and the recombinase Cre genes were inserted as independent reading frames between the virus genes N and P, flanked by the duplicated N/P intergenic region. A NotI restriction site was inserted after the eGFP gene, as well as 4 nucleotides after the Cre gene to ensure the whole genome would meet the paramyxovirus “rule of six”. Three helper plasmids were made by cloning NP, P, and L genes of SeV Cantell into the pTM1 vector. All these plasmids were confirmed by nanopore sequencing.

To rescue the recombinant virus, BSR-T7 cells grown in DMEM containing 10% FBS, 50ng/mL Gentamicin, 1mM Sodium Pyruvate, and 2mM L-Glutamine were transfected with a plasmid’s mixture containing 4.0 μg pSL1180-rSeV-C^eGFP-Cre^ 1.44 μg pTM1-NP, 0.77 μg pTM1-P and 0.07 μg pTM1-L using Lipofectamine LTX according to manufacturer’s guidelines. After a 5 h incubation, the medium was changed to infection medium (DMEM containing Pen/Strep, 35% Bovine Serum Albumin (BSA) (Sigma), 5% NaHCO_3_) with 1 μg/mL TPCK-treated trypsin (Worthington Biochem. Corporation), then cells were incubated at 37°C. The monolayers were monitored daily for eGFP expression and harvested on day 4 post-transfection. After 3 freeze-thaw cycles, the supernatants were clarified by centrifugation and used to infect 10-day-old embryonated chicken eggs through the allantoic cavity. After incubation for 40 hours at 37°C, 40 – 70% humidity, the allantoic fluids were harvested and the TCID_50_ was measured using LLC-MK2 cells.

### RNA extraction and RT-qPCR

Lung samples were homogenized in TRIzol (Ambion Inc.), and total RNA was extracted following manufacturer’s guidelines. To remove any trace of DNA contaminants, 1 µg of total RNA was treated with DNAse I (Thermo scientific) following manufacturer’s guidelines and cDNA synthesis was then carried out with the High-Capacity cDNA Reverse Transcription kit (Applied biosystems). For quantitative analysis by RT-PCR (qPCR), 10 ng/µL of cDNA was amplified using SYBR Green Mastermix (Thermofisher) in a BioRad C1000 Touch thermal cycler (BioRad). SeV NP (forward 5-TGCCCTGGAAGATGAGTTAG-3’, reverse 5’-GCCTGTTGGTTTGTGGTAAG-3’) relative copy numbers were normalized to mouse GAPDH (forward 5-CTCCCACTCTTCCACCTTCG-3’, reverse 5’-CCACCACCCTGTTGCTGTAG-3’) and corrected for mouse alpha-Tubulin (forward 5’-TGCCTTTGTGCACTGGTATG-3’, reverse 5’-CTGGAGCAGTTTGACGACAC-3’) expression as described previously^22^.

### Generation of recombinant huFc-SeV NP antibody

Sequence of the mouse variable heavy and kappa chains were obtained by using SMARTer 5’ RACE technology (Takara Bio, USA) adapted for antibodies to amplify the variable genes from heavy and kappa chains for each hybridoma based on isotype. Briefly, RNA was extracted from each hybridoma using Qiagen RNeasy Mini Kit (Qiagen, Valencia, CA), followed by first stand cDNA synthesis using constant gene specific 3’ primers (GSP) based on the specific isotype of the hybridoma and incubation with the SMARTer II A Oligonucleotide and SMARTscribe reverse transcriptrase. Amplifying PCR of the first stand cDNA product was then performed using SeqAmp DNA Polymerase (Takara) with a nested 3’ primer to the constant genes and a 5’ universal primer based on universal primer sites added to the 5’ end during cDNA generation. Purified PCR product was then submitted for Sanger sequencing using 3’ constant gene primers (GeneWiz, South Plainfield, NJ). Sequence results were blasted against the IMGT human databank of germline genes using V-Quest (http://imgt.org) and analyzed for CDR3/junction identity and V(D)J usage. Clones were chosen from each clonal family, and DNA was synthesized and cloned in-framed into pcDNA3.4 vector containing a human IgG1 constant region and a human kappa light chain constant region, making a chimeric mouse variable/human constant antibody. (GenScript USA Inc., Piscataway, NJ). Heavy and light cloned plasmids were mixed 1:1 and transfected into Expi293 cells according to manufacture protocol (ThermoFisher). Supernatants were harvested five days later and purified on protein A/G Hi-Trap columns on an AKTA FPLC (Cytiva). Antibody was eluted off the columns at low pH and dialyzed against 1xPBS. Antibody quantitation was performed by 280/260 absorbance on a Nanodrop spectrophotometer (DeNovix).

### Histopathology and tissue immunofluorescence

Mice lungs were perfused with 8 mL PBS and inflated with 0.7 mL of OCT compound (Tissue-Tek) mixed 1:1 v/v with 4% paraformaldehyde (Electron Microscopy Sciences) diluted in PBS. Inflated lungs were snap-frozen and stored at -80°C until sectioning. Tissue sections (4 µm) were stained with hematoxylin and eosin, and chronic lung disease was scored on a scale of 0 to 3 for alveolitis, peribroncholitis and airway metaplasia. Percentage values and area affected was determined, multiplied by the intensity scores previously defined, and the resulting weighted scores were graphed. For immunostaining analysis, tissue sections were washed in PBS to remove OCT, and Fc receptor blockade was performed using anti-mouse FcγRIII/II (FcBlock) (Jackson Immunoresearch) diluted 1:200 in PBS containing 1% bovine serum albumin (BSA). Surface staining was performed using a panel of antibodies targeting T cells, B cells, macrophages, and dendritic cells (**Supplementary Table 1**) at 4°C overnight. Sections were washed in PBS and surface antibodies were detected with anti-Rat AlexaFluor-488 secondary antibody (BioLegend). After surface staining, sections were permeabilized with 0.2% Saponin (Sigma) diluted in PBS containing 1% BSA and FcBlock (1:200), and intracellular staining was performed using recombinant hu-mouse anti SeV-NP antibody conjugated with AlexaFluor-647 (1:1000) (Invitrogen) in combination with either anti-Krt5 (1:500) (BioLegend) or anti-Krt8 (1:200) for 1 h at room temperature. Intracellular primary antibodies were detected with anti-Rabbit AlexaFluor 488 or anti-Rat AlexaFluor 488 secondary antibodies. Nuclear staining was performed with Hoechst (Invitrogen) and tissue autofluorescence was quenched using 1 x True Black (Biotium) diluted in 70% ethanol. Slides were mounted with Fluormount-G (Invitrogen) and images were acquired using a Zeiss Axio observer Widefield fluorescence microscope, using 5x and 20x objectives.

### Fluorescence RNA *in situ* hybridization

RNA *in situ* hybridization was performed using RNAscope Multiplex Fluorescent Reagent Kit v2 (Advanced Cell Diagnostics, Inc., Newark, NJ, USA) according to the manufacturer’s instructions. Lung sections (4 μm) were washed as described in the previous paragraph and hybridized for 2 h at 40°C with the RNAscope probe V-SeV-NP-C1 (Cat. No. 1118511-C1) targeting the genomic RNA sequence of NP gene. Preamplifier, amplifier, HRP-labeled oligos, and TSA plus (Cyanine3 or Cyanine5) (Akoya biosciences) dye was then hybridized at 40°C. Nuclear staining was performed with DAPI, and images were acquired as described in the previous sub-session.

### Multicolor flow cytometry and cell sorting

At 3 and 49 dpi, mouse lungs were inflated with 0.7 mL of digestion mix containing collagenase A (Sigma), dispase (Thermofisher), liberase TL (Sigma) and DNAse I (Sigma) and incubated at 37°C for 30 min with agitation. Digested samples were then briefly vortexed and filtered through a 70-µm filter mesh to obtain single-cell suspensions. The obtained cells were washed with PBS containing 5% FBS, treated with red blood cells lysis buffer (Sigma), and total viable cells were quantified with trypan blue staining using an automated cell counter (TC-20 Automated Cell Counter; BioRad). For each sample, 2×10^6^ cells were resuspended in PBS supplemented with 1% BSA and 2 mM EDTA (Corning). Next, Fc receptor blockade and viability staining were simultaneously performed using Rat anti-mouse FcγRIII/II (CD16/32; BD Biosciences) and ZombieNIR (BioLegend) for 10 min at room temperature. To define major cell subpopulations (**Fig S1**) we employed a panel of 16 antibodies (**Table S1**) to stain for surface markers. For intracellular staining of Sendai virus NP, cells were fixed/permeabilized with the FoxP3/ Transcription Factor Staining Buffer Set (eBioscience) following manufacturer’s guidelines, and incubated with recombinant hu-mouse anti-SeV NP antibody, conjugated with AlexaFluor-647 (Invitrogen) for 1h at 4°C.

To isolate specific lung cell populations with fluorescent activated cell sorting (FACS), surface staining was performed as mentioned above using a panel of lineage-specific markers (**Supplementary Table 1**). Live CD45^+^, Lineage^-^, Thy1.2^+^ and sm13^+^ cells were defined as ILC2s. Macrophages were obtained from the same single-cell suspensions mentioned above using the Anti-F4/80 MicroBeads UltraPure, mouse (Miltenyi Biotec) isolation kit. All flow cytometry experiments were performed using a Cytek Aurora spectral flow cytometer (Cytek Biosciences), with acquisition of at least 1×10^6^ total events. FACS experiments were performed using a BD-FACS Aria-II. Data analysis was done using FlowJo V12 software (Tree Star Inc.).

### RNA-seq of sorted cells

For both ILC2 and macrophages obtained from SeV-infected sm13 mice, and for tdTom^+^, tdTom^+^NP^+^, and negative cells obtained from rSeV-C^eGFP-Cre^-infected tdTom mice, total RNA was extracted from at least three cell pools per condition, using KingFisher APEX automated RNA Extraction and Purification system (ThermoFisher) according to the manufacturer’s guidelines. Each ILC2 and Macrophage sample, as well as tdTom^+^, tdTom^+^NP^+^, and negative cell samples was a resulting pool of cells from 2 individual animals. Total cDNA libraries were prepared from 100 ng of starting RNA using TruSeq Total RNA Library Prep Kit, with subsequent Ribo-Zero Human/Mouse/Rat Sample Prep Kit following manufacturer’s instructions. Libraries were run on Illumina NovaSeq 6000 to generate 150 bp, paired-end reads, resulting in 79-120 million reads per sample with an average Phred score of 35.75.

### Viral reads and host transcriptome analysis

Sequencing adaptors were removed from the raw sequencing data using Cutadapt^49^. Trimmed reads were then mapped to the mouse transcriptome (Ensembl release 79, EnsDb.Mmusculus.v79) using Kallisto, with 60 boostraps per sample^50^. Subsequent import and annotation of transcripts were done in R environment using the TxImport package^51^. Differentially expressed genes (DEGs) (p value < 0.05, fold change > 2) between pairwise comparisons were obtained by linear modeling and Bayesian statistics using the VOOM function from the Limma package^52^. Gene Ontology (GO) analysis was done using the enrichGO function from the ClusterProfiler package^53^ with a p value cut-off of 0.05. Gene Set Enrichment Analysis (GSEA) was performed using the Molecular Signatures Database (MSigDB) msigdbr R package^54^ including pathways found in the C2, C5 and H Mus Musculus gene collections^55,56^. Finally, reads mapping the mouse genome were removed from the trimmed dataset using Bowtie2 v2.4.1^57^ and virus coverage was obtained using SAMtools v1.15^58^ after aligning the non-host reads to the rSeV-C^eGFP-Cre^ genome using Bowtie2. Virus coverage per sample was visualized using the ggplot2 R package^59^. All data analysis was performed in RStudio (v. 2023.06.0+421).

### Statistical analysis

Statistical significance was inferred using GraphPad Prism software version 9.0 (GraphPad Software, San Diego, CA). For animal experiments, group size consisted of 3-7 mice per group. The weight-loss curve was analyzed by calculating the area under the curve (AUC) from both groups and comparing them using student t-tests. One-way and Two-way analysis of variance (ANOVA) with either Holm-Sídák or Bonferroni post-test was used to estimate the statistical significance between conditions of the remaining experiments. P values <0.05 were considered significant.

### Data deposition

Next generation sequencing raw data of SeV-52 and rSeV-C^eGFP-Cre^ experiments described in Figs 3, 4, and 5 was deposited in SRA under accession number PRJNA1034107.

## Acknowledgements

We thank J. Andrew Duty from the Center for Therapeutic Antibody Development, Drug Discovery Institute, Icahn School of Medicine, Mount Sinai, New York, NY, USA for the technical support in the generation of the SeV humanized antibody used in this study. We thank the Tissue Histology Core at the Center for Reproductive Health Sciences, Department of Obstetrics & Gynecology, Washington University School of Medicine, Saint Louis, MO, USA for all the support with the tissue sectioning and histologic staining. We also thank the Flow Cytometry & Fluorescent Activated Cell Sorting Core at the Department of Pathology & Immunology, Washington University School of Medicine, Saint Louis, MO, USA for the help with the cell sorting experiments.

This project was funded by NIH grants AI127832 and A137062, and the Washington University BJC Investigator program (to C.B.L), NIH grants HL148033, AI176660, and AI163640 (to S.J.V.D.), and the National Research Korea Award NRF-2020 R1A6A3A03037855 (to D.-H.K).

## Declaration of interests

The authors declare no competing interests.

---

**Figure S1.**
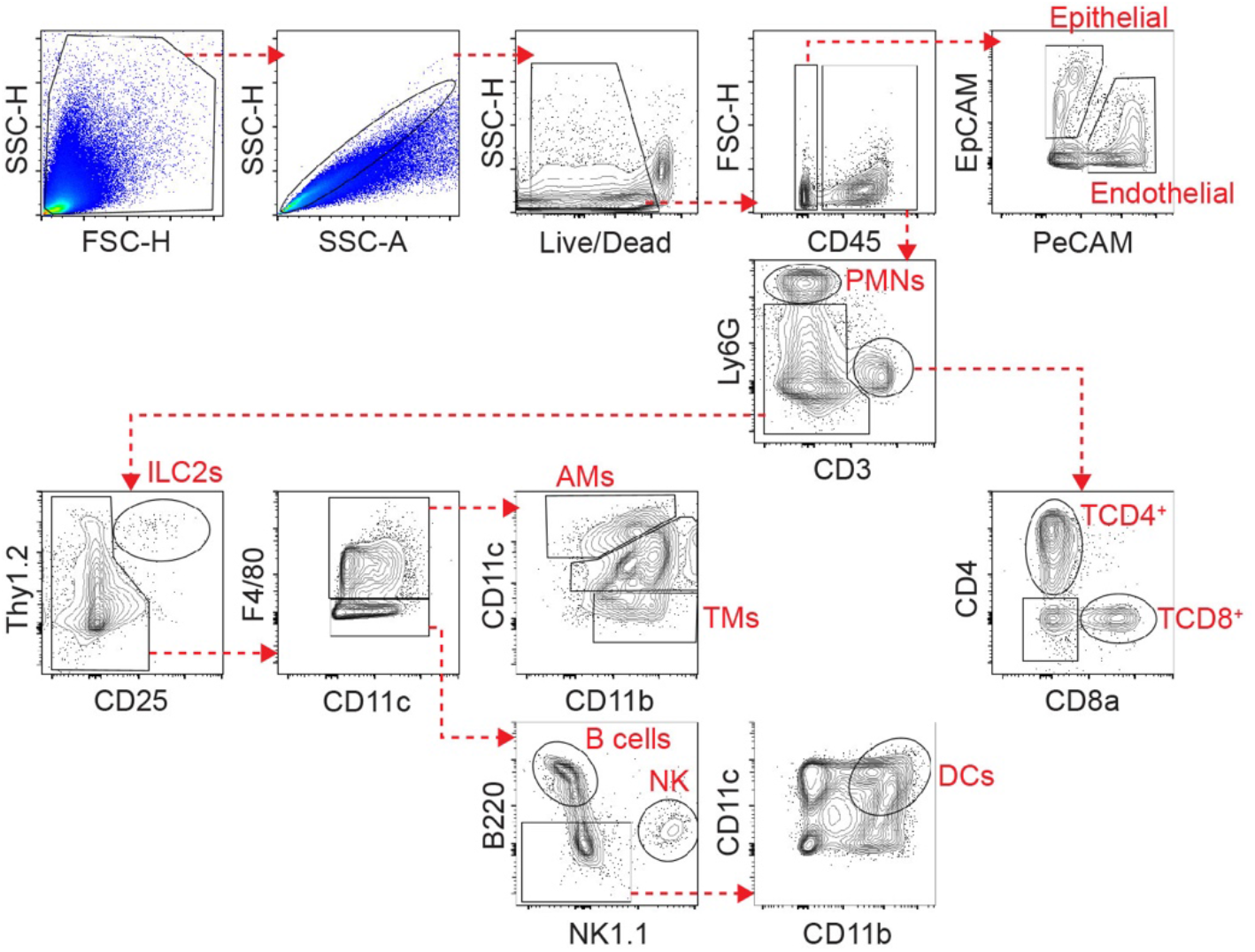
Gating strategy for spectral flow cytometry analysis of SeV-infected lungs. Representative dot plots indicating the following cell subsets defined from live cells: Epithelial cells (CD45^-^EpCAM^+^PeCAM^-^), Endothelial cells (CD45^-^EpCAM^-^PeCAM^+^), Polymorphonuclear cells (PMNs) (CD45^+^Ly6G^+^CD3^-^), T CD4^+^ lymphocytes (CD45^+^Ly6G^-^CD3^+^CD4^+^CD8a^-^), T CD8^+^ lymphocytes (CD45^+^Ly6G^-^CD3^+^CD4^-^CD8a^+^), Type 2 innate lymphoid cells (ILC2s) (CD45^+^Ly6G^-^ CD3^-^Thy1.2^+^CD25^+^), Alveolar macrophages (AMs) (CD45^+^Ly6G^-^CD3^-^F4/80^+^CD11c^hi^CD11b^low^), Tissue macrophages (TMs) (CD45^+^Ly6G^-^CD3^-^F4/80^+^CD11c^low^CD11b^hi^), B cells (CD45^+^Ly6G^-^ CD3^-^F4/80^-^B220^hi^), Natural killer cells (NK) (CD45^+^Ly6G^-^CD3^-^F4/80^-^B220^-^NK1.1^hi^), Dendritic cells (DCs) (CD45^+^Ly6G^-^CD3^-^F4/80^-^B220^-^NK1.1^-^CD11c^hi^CD11b^hi^).

**Figure S2.**
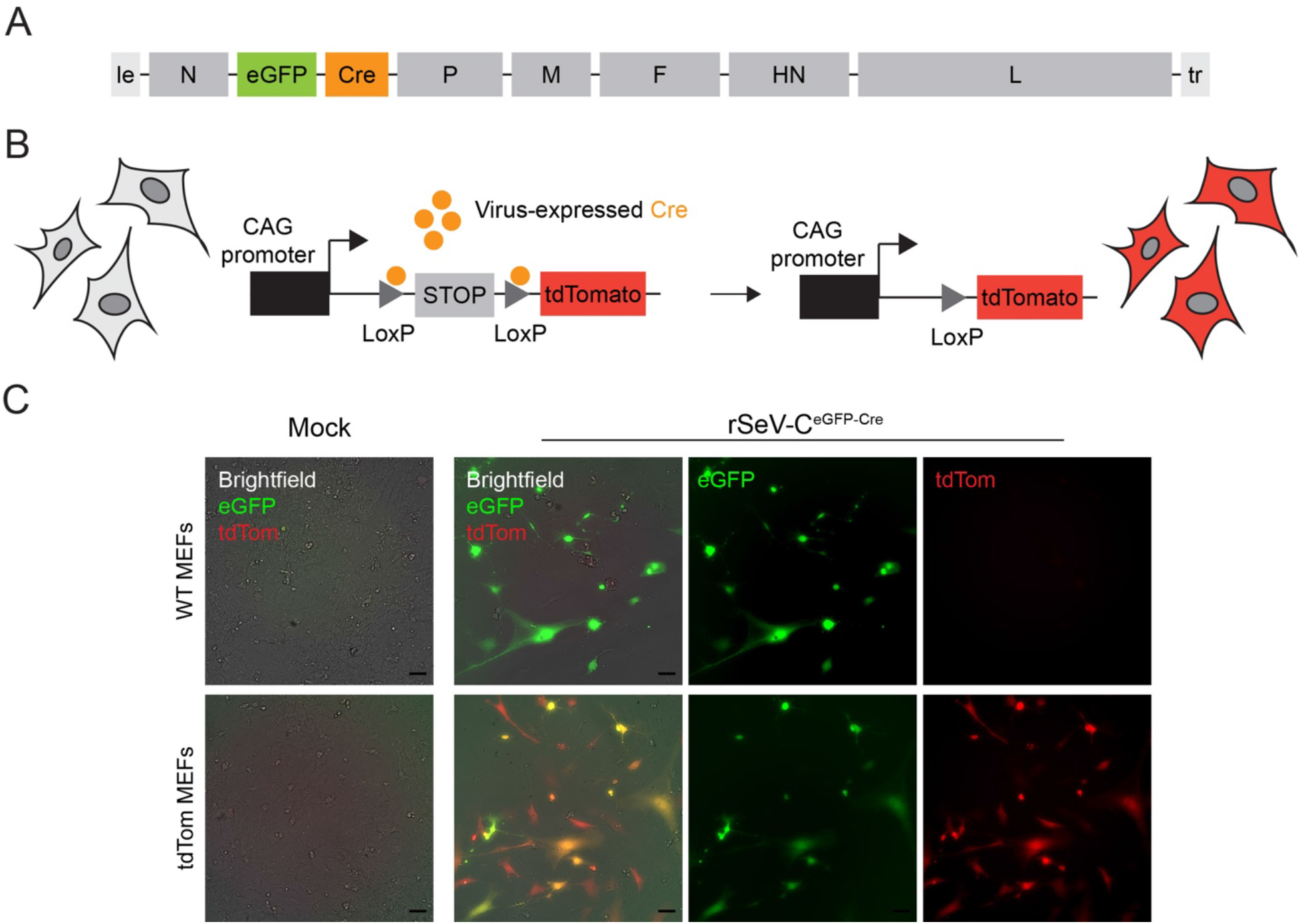
Generation of a Cre-expressing recombinant Sendai virus. **A.** Genome schematics showing the insertion of eGFP and the recombinase Cre genes as independent reading frames in the SeV Cantell genome, between the virus genes N and P, to generate a Cre-expressing SeV recombinant virus (rSeV-C^eGFPCre^). **B.** Schematic design showing Cre recombination of the LoxP-STOP-tdTomato reporter gene cassette leading to constitutive expression of the tdTomato fluorescent protein. **C.** Murine embryonic fibroblasts (MEFs) from either WT C57BL/6 mice or tdTomato reporter mice were infected *in vitro* at a multiplicity of infection (MOI) of 0.01 TCID_50_/cell to test the robustness of the reporter system. Representative images from two independent experiments were taken after 24 hpi using a widefield fluorescence microscope. eGFP signal (green) and tdTomato signal (red) were overlaid on brightfield images of WT and tdTomato-infected MEFs. 20x magnification. Scale bars: 50 µm.

**Figure S3.**
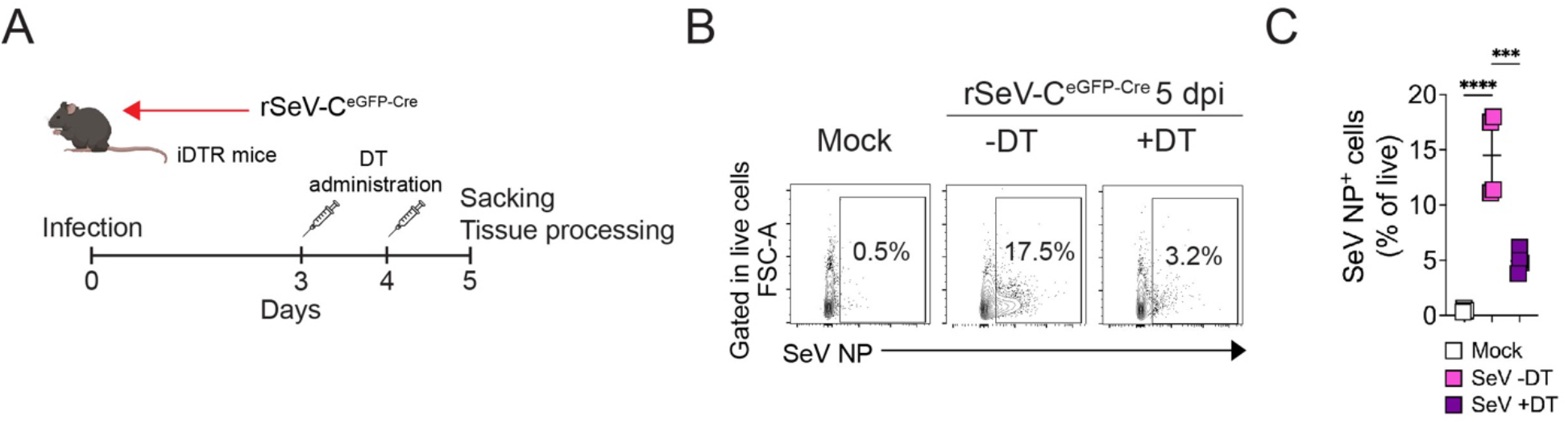
Diphtheria toxin administration efficiently depletes SeV infected cells. **A.** Experimental design showing diphtheria toxin (DT) regime treatment and timepoints for tissue harvesting and analysis. **B.** Representative dot plots of rSeV-C ^eGFP-Cre^ -infected lungs analyzed by flow cytometry for SeV NP expression. 50,000 events acquired. **C.** Quantification of SeV NP^+^ cell frequency (% of live) detected by flow cytometry at 5 dpi. Mean values ±SD are displayed, data are representative of 2 individual experiments, 4-5 mice per group. One way ANOVA with Bonferroni post-test was used to estimate statistical significance between groups. ***P<0.0005, ****P<0.0001.

**Table S1.** Key reagents.

| Reagent | Company | Catalog N. |
| --- | --- | --- |
| **Antibodies** |  |  |
| Anti-mouse EpCAM (CD326) BV421 antibody | Biolegend | 118225 |
| Anti-mouse Ly6G eFluor450 antibody | eBioscience | 48-5931-82 |
| Anti-mouse CD8a BV570 antibody | Biolegend | 100740 |
| Anti-mouse PeCAM (CD31) BV605 antibody | Biolegend | 102427 |
| Anti-mouse CD4 BV711 antibody | Biolegend | 100447 |
| Anti-mouse F4/80 BV785 antibody | Biolegend | 123141 |
| Anti-mouse Thy1.2 (CD90.2) AlexaFluor 488 antibody | Biolegend | 105316 |
| Anti-mouse NK1.1 BB700 antibody | BD Biosciences | 566502 |
| Anti-mouse B220 PerCP-eFluor 710 antibody | eBioscience | 46-0452-82 |
| Anti-mouse CD25 PE antibody | eBioscience | 12-0251-81 |
| Anti-mouse CD64 PE-Cy5 antibody | ThermoFisher | MA5-38711 |
| Anti-mouse CD11c PE-Cy7 antibody | eBioscience | 25-0114-82 |
| Anti-mouse CD11b AlexaFluor 700 antibody | BD Pharmingen | 557960 |
| Anti-mouse CD45.2 APC-eFluor 780 antibody | eBioscience | 47-0454-82 |
| Anti-mouse CD3 APC-Fire 810 antibody | Biolegend | 100268 |
| Anti-mouse CD11c FITC antibody | eBioscience | 11-0114-85 |
| Rabbit anti-mouse Krt5 antibody | Biolegend | 905504 |
| Rat anti-mouse Krt8 antibody | DSHB | TROMA-I |
| Rat anti-mouse CD3 antibody | BD Biosciences | 555273 |
| Rat anti-mouse F4/80 antibody | BD Biosciences | 565409 |
| Rat anti-mouse B220 antibody | BD Biosciences | 553084 |
| Rat anti-mouse Thy1.2 (CD90.2) antibody | Biolegend | 140301 |
| Rat anti-mouse CD16/CD32 (FcBlock) | Biolegend | 101320 |
| anti-mouse CD3 biotin-conjugated | Biolegend | 309806 |
| anti-mouse CD4 biotin-conjugated | Biolegend | 100244 |
| anti-mouse CD8a biotin-conjugated | Biolegend | 100508 |
| anti-mouse/human CD11b biotin-conjugated | Biolegend | 11704 |
| anti-mouse Ly6G/Ly6C (Gr-1) biotin-conjugated | Biolegend | 101204 |
| anti-mouse NK1.1 biotin-conjugated | Biolegend | 108403 |
| anti-mouse CD49b (Pan-NK) biotin-conjugated | Biolegend | 108704 |
| anti-mouse CD19 biotin-conjugated | Biolegend | 108903 |
| anti-mouse Ter-119 (Erythroid cells) biotin-conjugated | Biolegend | 115504 |
| anti-mouse CD11c biotin-conjugated | Biolegend | 116203 |
| anti-mouse TCR gamma/delta biotin-conjugated | Biolegend | 117304 |
| anti-mouse CD170 (Siglec-F) biotin-conjugated | Biolegend | 118103 |
| anti-mouse CD90.2 (Thy1.2) Antibody biotin-conjugated | Biolegend | 155512 |
| anti-human CD4 Antibody biotin-conjugated | Biolegend | 140314 |
| Goat Anti-Rabbit AlexaFluor488 Secondary antibody | Invitrogen | A-11008 |
| Anti-Rat AlexaFluor488 Secondary antibody | Biolegend | 405418 |
| Goat Anti-Rat AlexaFluor647 Secondary antibody | Invitrogen | A-21247 |
| **Commercial assays** |  |  |
| RNAscope® Multiplex Fluorescent Detection Kit v2 | ACD | 323110 |
| RNAscope® Target Retrieval Reagents | ACD | 322000 |
| RNAscope® 2.5 HD Detection Reagents-BROWN | ACD | 322310 |
| TSA Plus Fluorescein Kit 50-150 slides | Akoya biosciences | NEL741001KT |
| TSA Plus Cyanine 3 Kit 50-150 slides | Akoya biosciences | NEL744001KT |
| TSA Plus Cyanine 5 Kit 50-150 slides | Akoya biosciences | NEL745001KT |
| Power SYBR™ Green PCR Master Mix | ThermoFisher | 4367660 |
| High Capacity cDNA Reverse Transcription Kit | Applied Biosystems | 4368813 |
| Zombie NIR™ Fixable Viability Kit | Biolegend | 423106 |
| AlexaFluor 647 Antibody Labeling kit | ThermoFisher | A20186 |
| **General reagents** |  |  |
| BSA | Sigma | A3059-100G |
| TPCK-treated Trypsin | Worthington Biochem. Corporation | #LS003750 |
| EDTA | Corning | 46-034-CI |
| DNase I (for molecular biology) | ThermoScientific | #EN0252 |
| Paraformaldehyde (PFA) | Electron Microscopy Sciences | 15714-5 |
| Saponin | Sigma | 47036-50G-F |
| Collagenase A | Sigma | 10103586001 |
| Dispase | Gibco | 17105-041 |
| Liberase TL | Sigma | 5401020001 |
| DNase I (for tissue digestion) | Sigma | 10104159001 |

## Notes

### Competing Interest Statement

The authors have declared no competing interest.

